# Encoding the expectation of a sensory stimulus

**DOI:** 10.1101/281071

**Authors:** Lijun Zhang, Alex Chen, Debajit Saha, Chao Li, Baranidharan Raman

**Author notes:** These authors contributed equally to this manuscript (Lijun Zhang, Alex Chen, Debajit Saha).

## Abstract

Most organisms possess an ability to differentiate unexpected or surprising sensory stimuli from those that are repeatedly encountered. How is this sensory computation performed? We examined this issue in the locust olfactory system. We found that odor-evoked responses in the antennal lobe (downstream to sensory neurons) systematically reduced upon repeated encounters of a temporally discontinuous stimulus. Rather than confounding information about stimulus identity and intensity, neural representations were optimized to encode equivalent stimulus-specific information with fewer spikes. Further, spontaneous activity of the antennal lobe network also changed systematically and became negatively correlated with the response elicited by the repetitive stimulus (i.e. ‘a negative image’). Notably, while response to the repetitive stimulus reduced, exposure to an unexpected/deviant cue generated undamped and even exaggerated spiking responses in several neurons. In sum, our results reveal how expectation regarding a stimulus is encoded in a neural circuit to allow response optimization and preferential filtering.

## Introduction

The ability to adapt is key to the survival of many living organisms^1^. While this computational task appears relatively straight-forward, any solution should satisfy at least two important constraints or requirements. First, attenuation of stimulus-evoked responses should not alter the identity and the intensity of the repetitive or familiar stimulus. This constraint arises due to findings that behavioral preferences for many stimuli vary with intensity^2-4^. Second, only the response evoked by the familiar stimulus should be selectively impeded, and sensitivity to novel or unexpected cues should not be compromised. In this study, we explored how the locust olfactory system addresses these issues.

In the olfactory system, adaptation occurs right at the level of olfactory receptor neurons. Usually, sensory neuron responses to prolonged chemical exposures reduce over the duration of that exposure^5-9^. However, a lengthy time-window of non-exposure to the stimulus, typically on the order of tens of seconds, can allow full recovery of the sensory neuron response strength^3,7,10^. Interestingly, such temporal discontinuity in stimulus encounters does not prevent the stimulus-evoked responses in downstream centers from diminishing upon subsequent encounters of the familiar stimulus^11-13^. This suggests that information about the temporally discontinuous but repetitive stimuli continue to persist even in the absence of olfactory sensory neuron input.

If sensory memory persists in the early olfactory circuits, how does it interfere with subsequent responses evoked by the familiar cue that caused this short-term memory? Stimulus-specific sensory adaptation can be achieved relatively easily in sensory systems with labeled-line coding schemes. However, it becomes particularly challenging in a modality such as olfaction where most chemosensory cues are encoded by spatiotemporal patterns of neural activities distributed across an ensemble of overlapping sets of neurons^14-19^. In the vertebrate and the invertebrate olfactory systems, each olfactory receptor neuron and their downstream targets (projection neurons in invertebrate antennal lobe or mitral/tufted cells in the vertebrate olfactory bulb) respond to multiple stimuli ^18-24^. Conversely, most odors activate an overlapping set of neurons in these early processing stages. To add further complexity, the set of neurons activated is not static but has been shown to evolve over time. Given this dynamic and combinatorial nature of odor-evoked neural representations, how are neural responses to a repetitive olfactory stimulus altered? Do alterations of neural response strength upon repetition confound encoding of stimulus intensity? More importantly, how does adaptation to a stimulus alter processing of other cues? We explored these issues in this study using a model of the invertebrate olfactory system.

## Results

### Sensory memory of a temporally discontinuous stimulus

We began by looking at how populations of olfactory sensory neurons and their followers in the antennal lobe adapted their responses to a repeatedly encountered odorant (**Fig. 1a-b**; 4 s long odor pulses separated by a 60 s inter-pulse interval). Our results indicate that the sensory neuron inputs remain stable across multiple repetitions of the same odorant. No adaptation to an odorant was observed when the separation between repeats was on the order of tens of seconds. However, projection neuron (PN) responses in the downstream antennal lobe substantially reduced over repetitions of such temporally discontinuous stimulus. The PN response attenuated after the first encounter of the odorant. These observations suggest that the antennal lobe neural network must carry some information about the repeated odorant even in the absence of continuous sensory input.

**Fig. 1:**
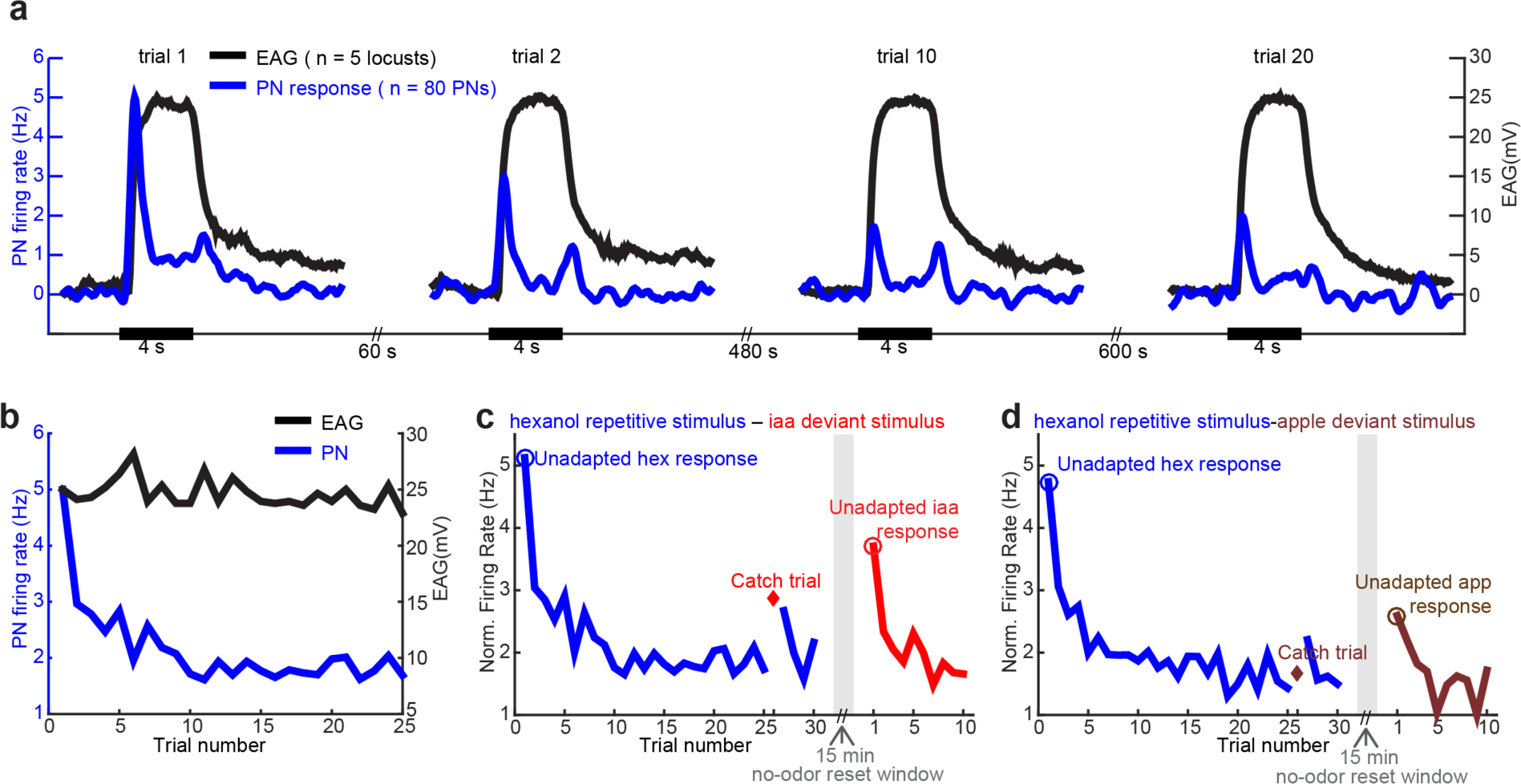
Neural responses to repeated temporally discontinuous stimulus. **(a)** Electroantennogram (EAG) measurements for four representative trials are shown in black. Signals were baseline aligned and normalized to the peak response of the first trial. Blue traces show population-averaged projection neuron (PN) firing rates over time for the corresponding trials. PN spike rates were also baseline aligned and normalized to the peak response observed during the first trial. The black bar along the x-axis represents odor presentation window (4 s). Note that between odor presentations in consecutive trials, there is a 60 second inter-stimulus period. **(b)** Peak odor-evoked EAG (black) and population PN firing rate (blue) responses are shown as a function of trial number. **(c)** The peak population PN firing rate evoked by a stimulus is shown as a function of trial number. Hexanol was repetitively presented during all but the twenty-sixth trial (‘catch trial’) in the first 30 trial block. During the ‘catch trial,’ a deviant stimulus (isoamyl acetate; iaa) was presented. Following the 30-trial block, no stimulus was presented for 15 minutes to allow the antennal lobe circuits to reset. Subsequently, ten trials of isoamyl acetate (iaa) were presented after this reset period. **(d)** Similar plot as in **panel c,** but using apple (app; a complex odorant) as the deviant stimulus.

Next, we examined whether the response strength attenuation was limited to the repeating stimulus alone. To understand this, we used a simple ‘oddball’ paradigm. We repeatedly presented one odorant before introducing a ‘catch trial’ (26^th^ trial in a 30-trial block), when a ‘deviant’ stimulus was presented. Note that the catch trial happened after many exposures to the repeating stimulus. We found that odor-evoked PN responses in the antennal lobe tended to increase during the catch trial (**Fig. 1c-d**). However, the peak response intensity was still lower than the ‘unadapted’ response evoked by the same deviant stimulus. Note that the unadapted response to an odor is defined here as the response elicited in the first trial after a fifteen-minute no-odor ‘reset’ period.

In sum, these results indicate that the antennal lobe neural network is the first neural circuit in the olfactory pathway where adaptation to a discontinuous but repeating stimulus occurs. Further, adaptation to one stimulus appears to impact how other cues are processed within the antennal lobe.

### Trial-by-trial changes in odor-evoked responses

Next, we examined how odor-evoked responses in individual PNs changed across twenty-five repeated presentations of the same stimulus (**Fig. 2a; Supplementary Fig. 2a**). We found that the odor-evoked neural activity patterns across trials were not a constant but changed in a systematic manner for individual PNs. We found that inhibition duration (PN4, PN5), response latency (PN2, PN9), response intensity, (PN1, PN6) and reliability (PN4, PN10) all varied on a trial-by-trial basis.

**Fig. 2:**
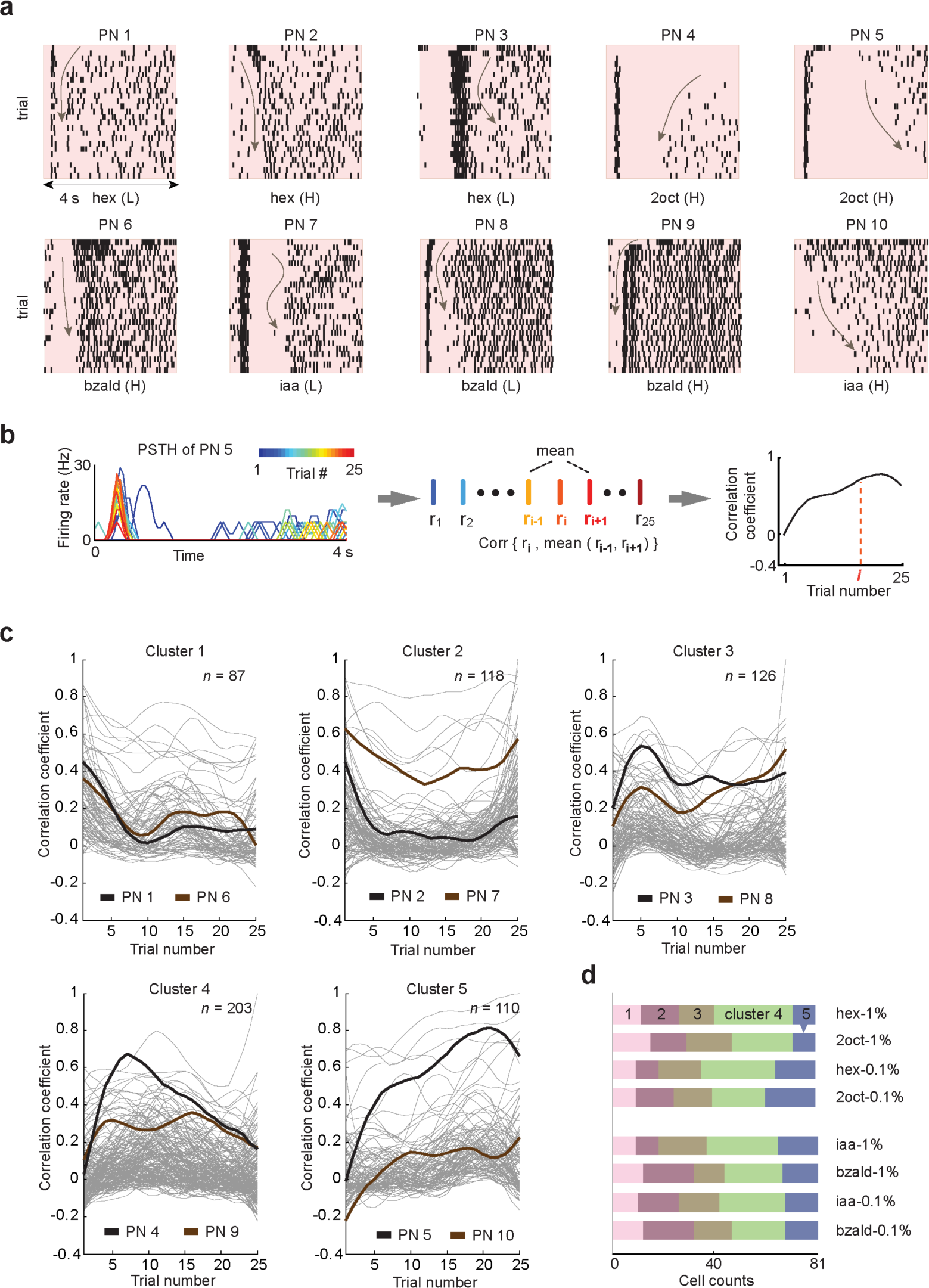
Inter-trial neural dynamics are diverse in individual projection neurons. **(a)** Responses of ten representative projection neurons (PNs) to various odorants are shown as raster plots (25 trials each). Arrows highlight the systematic changes in stimulus-evoked response features across trials: inhibition duration (PN1, PN4, PN5), response latency (PN2, PN9, PN10), response intensity (PN2, PN6) and reliability (PN3, PN7, PN8). Spiking activity during the entire four seconds of stimulus exposure is shown. **(b)** The neighboring correlation metric used to quantify the changes in spiking response patterns over trials is schematically shown. First, for each neuron, the stimulus-evoked spike trains elicited in a single trial were binned in non-overlapping 50 ms time windows (e.g. PN5, trial number is color coded). For a 4s odor exposure, this generated 80-dimensional vectors to characterize the PN response in each trial. The similarity between PN firing activity in a given trial and the average of its neighboring two trials (see **Methods**) was computed, and plotted as a function of trial number. **(c)** The neighboring correlation metrics for different PN-odor combinations clustered into five prominent response motifs (see **Methods**). After this categorization, evolution of the correlation coefficient as a function of trial number for each PN-odor combination is shown in five separate panels. Representative PNs, whose rasters are shown in **panel a**, are highlighted using darker colors. **(d)** Stacked bar plot summarizing the size of membership for each cluster is shown for each odorant used in this study. No significant differences between cluster membership distributions were found between stimuli (Two-way ANOVA, p>0.05).

To systematically quantify the trial-to-trial firing pattern changes, we employed a neighboring-trial similarity metric^25^ (**Fig. 2b**; see Methods). Briefly, to analyze each of the PN responses, we first counted the number of spikes in 50 ms non-overlapping time bins for each trial. As a result, the PN spiking activity to 4 s stimulus presentation in a given trial resulted in an 80-dimensional time series of spike counts. Then, we measured the similarity between responses observed in each trial with the mean responses observed in the trials immediately preceding and following (for the first and the last trial, we used two trials that immediately followed and preceded, respectively). These correlations were computed for each of the twenty-five trials and plotted as a function of trial number (**Fig. 2c**). Overall, this correlation metric resulted in various profiles for a total of 644 neuron-odor combinations.

To identify the predominant response motifs that characterize trial-to-trial response changes, we used an unsupervised clustering analysis. We found that neighborhood-correlation patterns for all PNs (all PN-odor combinations included) could be categorized into five major motifs or clusters (**Fig. 2d**; see Methods). Interestingly, we found that none of the response motifs suggested a monotonic increase (stable response patterns) or decrease (unstable response patterns) in the correlation metric over trials. However, in some subset of PNs, compared to the initial trials, there was an increase in correlation indicating stabilization of PN response patterns in later trials (e.g. cluster 5). In others, there was an overall decrease (e.g. cluster 1). Although the changes were more prominent in the first few trials, none of the clusters appeared to have fully converged onto steady response patterns even after twenty-five repeated exposures of the same odorant. We note that the same type of response variability was observed for all odorants and intensities tested, (Two-way ANOVA, p>0.05; **Fig. 2d**).

Therefore, we conclude that these systematic changes in PN responses indicate a non-converging, circuit-level adaptation triggered and sustained by repeated encounters of any discontinuous olfactory stimulus.

### Adaptation-invariant encoding of odor intensity

We next examined how the odor-evoked response strength varied across repeated trials of the same stimulus. Since the response strength also varies with stimulus intensity, we compared the changes due to reduction in intensity with those brought about by repetition. Our results indicate that the total spike counts (across all recorded PNs) reduced significantly within the first few trials for all odorants at all intensities. In general, the reduction was greater at higher intensities. For the two alcohols examined, the difference in spike counts between responses elicited at high and low stimulus intensities were maintained even after adaptation (**Fig. 3a**; two left-most panels). On the other hand, spike count differences between two different stimulus intensities diminished over trials for the other two odorants used (iaa and bzald; **Fig. 3a** two right-most panels). More importantly, the earlier trials of odor exposures at low intensities elicited a stronger response than those evoked during the later trials of the same stimulus but at a higher intensity. These results suggest that adaptation may potentially confound the representation of stimulus intensity.

**Fig. 3:**
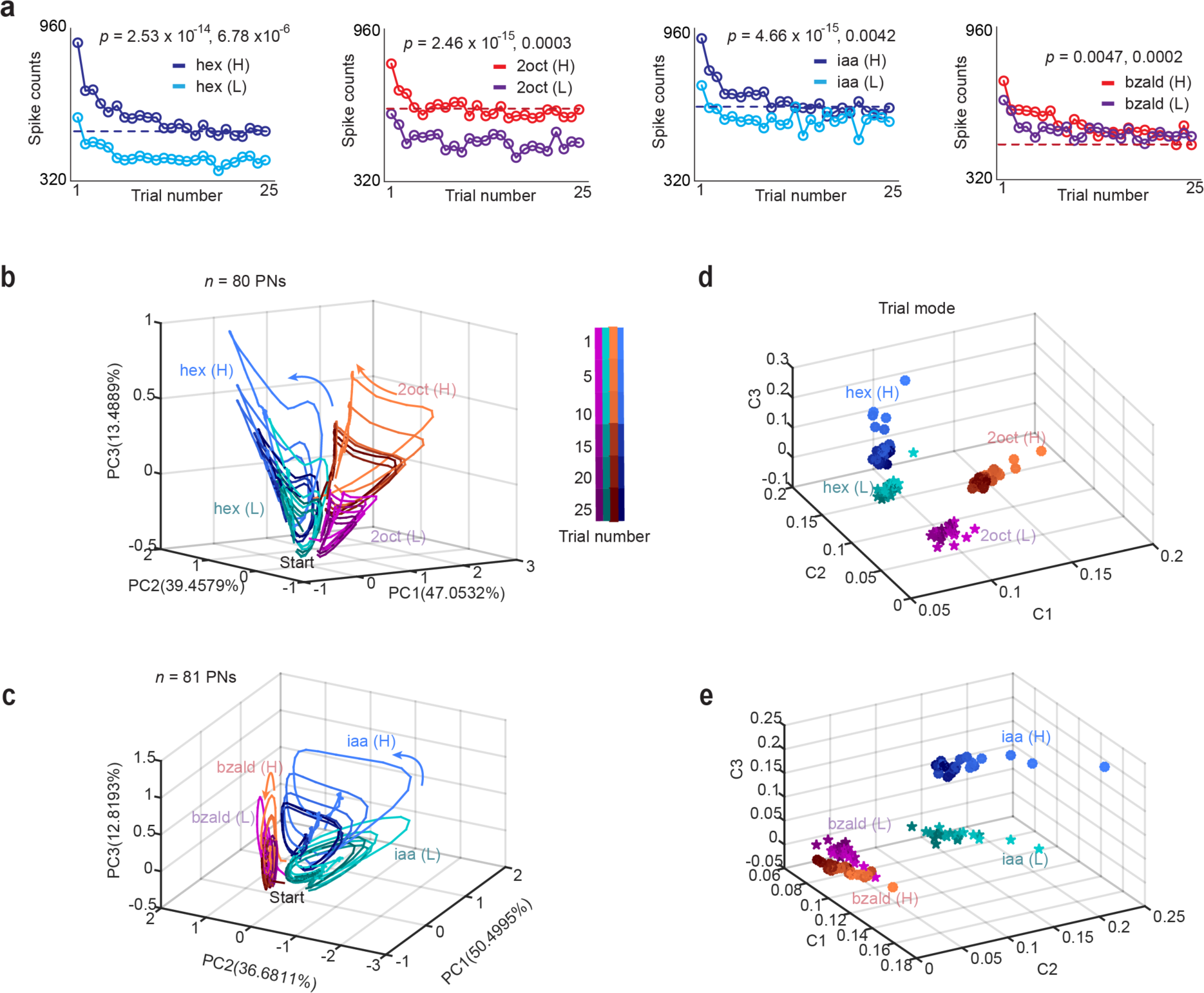
Ensemble neural activity change systematically over repeated trials. **(a)** Total spike counts across all PNs during the entire four seconds of stimulus exposure were calculated and plotted as a function of trial number. Each panel reveals the total odor-evoked response generated across all recorded PNs to two different intensities of the same stimulus (H = 1% by volume, L = 0.1% v/v). The dotted line indicates the minimum spike count observed during the 25^th^ trial of higher intensity odor presentations. Two-way ANOVA was used to compare the spike counts between different trials and different odorant concentrations (see **Methods**). **(b, c)** Odor-evoked ensemble projection neuron response trajectories are shown after dimensionality reduction using a tensor decomposition method (see **Methods**). For each odor pair, the region where the trajectories begin following odor onset (labeled ‘Start’), and the direction of response evolution over time are identified (colored arrows). The response trajectories of representative trials 1, 5, 10, 15, 20 and 25 are shown by color gradient from light (early trials) to dark (later trials). **(d, e)** Information retained (loading elements) in the trial dimension following a 3-rank tensor decomposition (see **Methods**) is plotted as a 3-d plot. Each trial is represented as a symbol in this plot. Color scheme same as that used in panels **b, c**. The solid circle and star symbols represent the higher (1%) and lower (0.1%) odorant concentrations, respectively.

Could the information regarding stimulus intensity be robustly encoded in the population neural responses? Previous studies have indeed shown the ensemble neural activities cluster based on odor identity and intensity^20,26^. However, this representation of stimulus-specific information stabilized only when the first trial responses were not considered. Given that major changes in spike counts due to adaptation occurred between the first and second encounters of a stimulus, we sought to examine how encoding of stimulus intensity varied before and after adaptation (i.e. across all trials).

To qualitatively understand this, we visualized the high-dimensional neural activities using a dimensionality reduction approach. Note that the spiking responses of many neurons (pooled across experiments) over time in each trial resulted in a 2-D matrix. The ensemble responses across multiple trials were concatenated to create a 3-D data cube (neuron x time x trial dimensions). Typically, this data cube is unfolded to a concatenated-matrix before performing dimensionality reduction^20,24^. Instead, we directly performed a 3-D tensor decomposition and approximate the data cube as a sum of three rank-one tensors (for a rank 3 approximation to facilitate visualization; see Methods). This pre-processing step followed by regular unfolding the data cube for linear dimensionality reduction, resulted in neural response trajectories that captured the trial-to-trial variations in the dataset better than the direct unfold-then-PCA approach^27,28^ (**Supplementary Fig. 1**).

We plotted the ensemble responses in each 50 ms time bin during the odor presentation time window (4 s), and linked them based on the order of their occurrence, to generate trial-by-trial odor response trajectories (**Fig. 3b, c)**. Note that each trial generated a single loop response trajectory after dimensionality reduction. Five such trajectories (shown in blue) correspond to the responses evoked by *hex* in representative trials: 1, 5, 10, 15, 20, and 25. Similarly, five *2oct* trajectories (red) for corresponding trials are also shown for comparison. Note that the response trajectories showed a systematic change from light colors (early trials) to darker colors (late trials). Notably, the population responses changed such that the trajectories evolved in similar directions, but the length of the trajectory monotonically reduced over repeated trials. Similar results were also observed for the other two odorants used in the study (**Fig. 3c**). Furthermore, we found that the basis vectors that best captured variance information in the trial dimension showed robust clustering of trials based on stimulus identity and intensity (**Fig. 3d, e; Supplementary Fig. 2b, c**). Hence, these results suggest that ensemble neural activity robustly encodes information about both odor intensity and identity even though the total number of spikes changes substantially with adaptation.

To quantify these observations, we summed the number of spikes over the entire odor exposure period (4 s) for each individual PN and compared the projection neurons response profiles generated by two different odorants or by the same odorant at two different intensities (**Fig. 4a-e**). We used a correlation-based distance metric to quantify the similarity between two population response profiles. We found that the combinatorial PN activity profiles varied substantially between odorants and subtly between different concentrations of the same odor, (**Fig. 4a-c**; *R* < 0.4, for hex vs. 2oct, and iaa vs. bzald *vs. R* = 0.692 for hex-1% vs. hex-0.1%; *R* = 0.467 for iaa-1% vs. iaa-0.1%). These results suggest that a combinatorial code can retain information about odor identity and intensity and is robust to changes that occur due to adaptation.

**Fig. 4:**
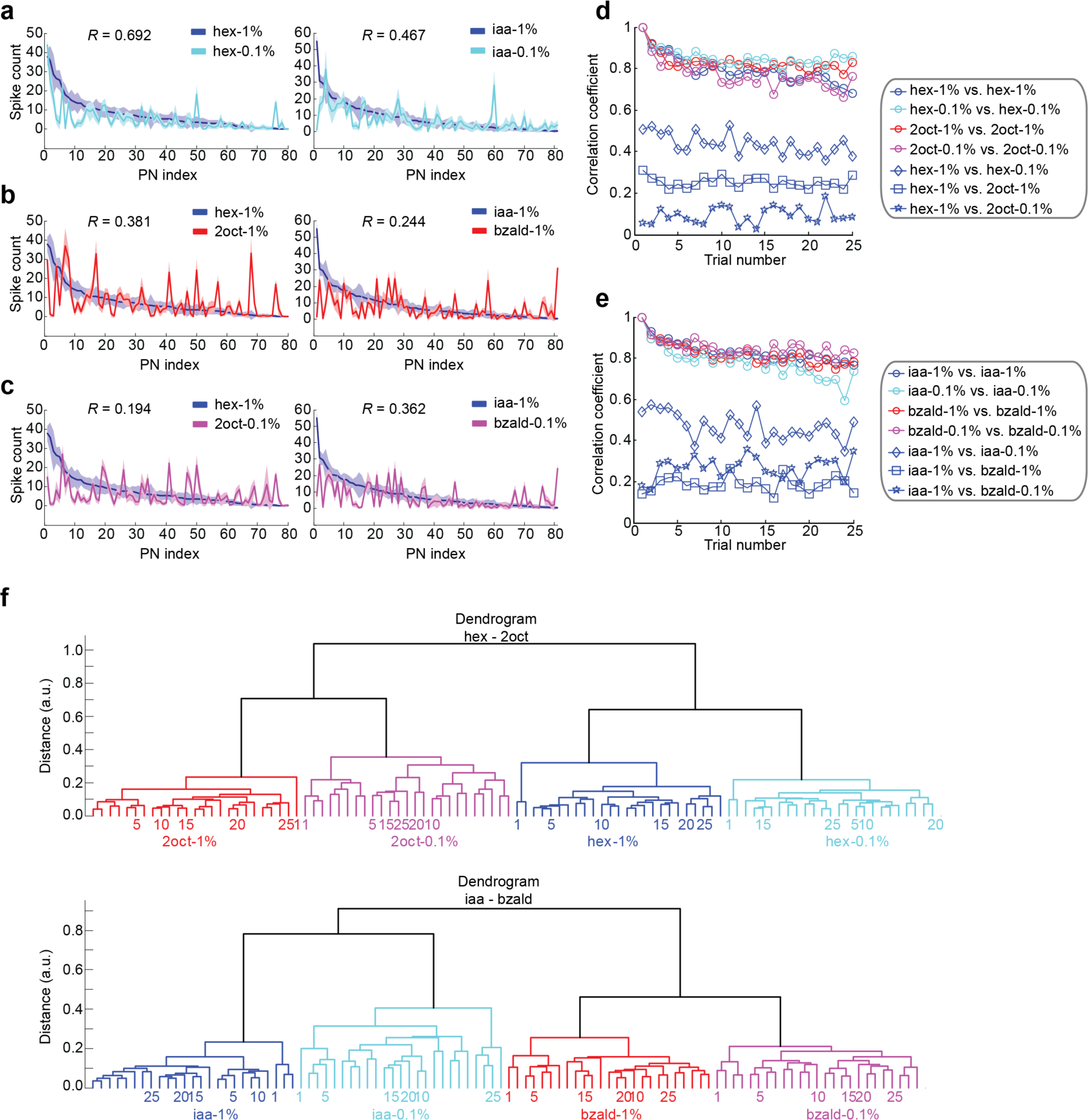
Odor identity and intensity information is maintained across trials. **(a)** Comparison of odor-evoked responses between high and low intensities of hexanol (left panel) and isoamyl acetate (right panel). Spiking activities of individual PNs were summed over the entire odor presentation window, sorted based on their response to the high-intensity presentations, and plotted to reveal the combinatorial response. Mean ± S.D. across the twenty-five trials are shown. Correlation coefficients (*R*) between the high vs low intensity PN response profiles for both odorants are shown in each panel. **(b)** Similar plots as in **panel a**, but comparing the PN response profiles evoked by different odorants presented at the same dilution level. **(c)** Similar plots as in **panel a**, but comparing the PN response profiles generated by different odorants presented at different dilution levels. **(d)** A comparison of combinatorial PN response profiles activated by the same odorant across trials (‘ ο’), same odorant across intensities (‘◊’), between different odorants presented at same intensity (‘□’) and between different stimuli presented at different intensity levels (‘★’) are shown as a function of trial number. Note that similarity with respect to the first trial response of that odorant is used for comparisons across trials. For comparisons between odorants and intensity levels, similarity with respect to the first trial of hexanol at 1% was computed and plotted. **(e)** Similar plot as in **panel d**, but analyzing responses of a different set of PNs to a different pair of odorants (iaa and bzald). For comparisons between odorants and intensities levels, similarity with respect to the first trial of isoamyl acetate at 1% was computed and plotted. **(f)** A dendrogram was generated using a correlation distance metric comparing trial-by-trial ensemble spiking activities evoked by two different stimuli at two different intensities (see **Methods**). Two major response clusters that correspond to stimulus identity and intensity were identified. The number at the leaf node represents the trial number.

Further, we examined the effect of adaptation on the combinatorial PN response profiles (**Fig. 4d, e**). We found that the correlation between response profiles changed systematically as a function of trial number (note that similarity between response profiles observed in a given trial with the first trial of the same stimulus is shown). However, these changes were subtle when compared with the changes due stimulus intensity and identity. Therefore, these results suggest that the stimulus information, including both odor identity and intensity, can be robustly encoded by the combinations of neurons activated even in the presence of neural adaptation.

Note that, the results in **Fig. 4d, e** were obtained based on comparison with the responses obtained in the very first trial. To generalize our conclusion, we performed a hierarchical clustering analysis of these combinatorial profiles (again a correlation distance metric was used, see Methods; **Fig. 4f**; Euclidean distance was used in **Supplementary Fig. 3** for comparison). We found that the ensemble response profiles were grouped first based on odor identity followed by intensity. Moreover, within each cluster of the same stimulus (indicated by the same color), the leaf nodes of the dendrogram tended to form sub-groups based on the trial number. Therefore, we conclude that the systematic changes within each stimulus condition only slightly perturbed but did not significantly alter the overall combinatorial profiles.

In sum, our results reveal that a combinatorial code could encode information regarding odor identity and intensity in an adaptation-invariant fashion. Since the same information about a stimulus is represented with fewer spikes in the later trials, we conclude that adaptation refines the odor codes by making them more efficient.

### Sensory memory creates a negative image of stimulus-evoked responses

Our results indicate that the antennal lobe response to a repeating stimulus can reduce even when the sensory inputs remain constant. This observation suggests that the sensory memory regarding previous encounters of the repeating stimuli must be maintained in the circuit even after its termination. We wondered whether the persistence of this memory could manifest in other ways. Therefore, we also examined how PN spontaneous activities changed over trials (**Fig. 5a-c**). Note that we analyzed spontaneous activities in 15 s time windows that preceded the odor presentation in each of 25 repeated trials of an odorant (**Fig. 5b**). We found that repeated exposure to an odorant altered spontaneous activity of individual neurons in an odor specific manner. Nearly two-thirds of the recorded PNs had significant differences in their spontaneous firing rates depending on the repeating odorant (**Fig. 5c)**. On the other hand, we found that spontaneous activities during two different epochs when the same stimulus was repeated were comparable (**Fig. 5b**). Further when the mean spontaneous firing rate across PNs in a given trial was visualized using principal component analysis, we found that each odorant at a given intensity formed a distinct cluster of baseline activities (**Fig. 5d; Supplementary Fig. 4**). Hence, these results indicate non-random, odor-specific changes in spontaneous PN firings when an odorant is repeatedly encountered.

**Fig. 5:**
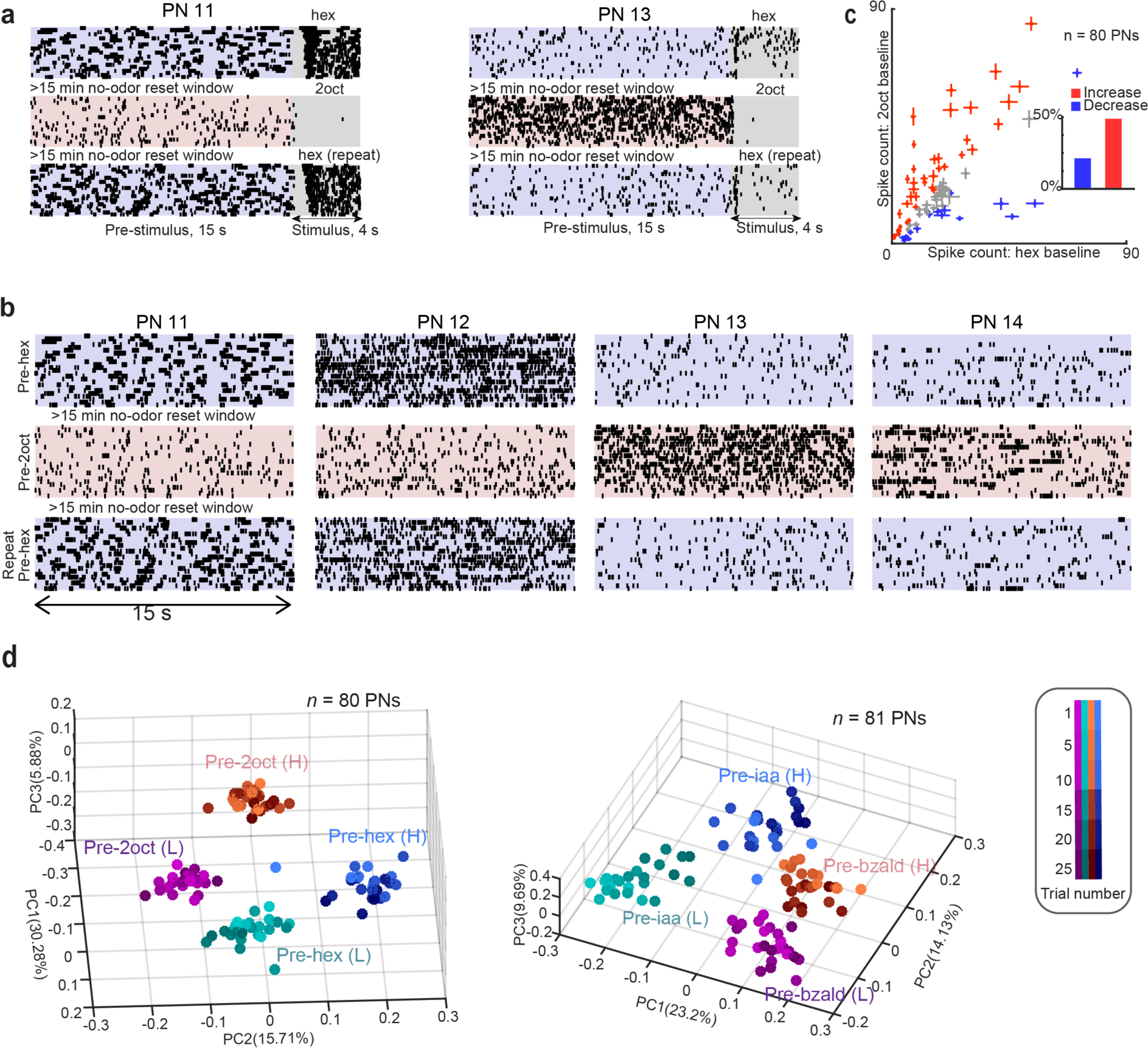
Sensory memory regarding the repeated stimulus persists even in the absence of that stimulus. **(a)** Raster plots revealing both pre-stimulus (15 s) and stimulus-evoked activities (4 s) of two representative projection neurons are shown. Each row corresponds to one trial, and spiking activity during twenty-five trials (top-most row – trial 1, bottom-most row – trial 25) are shown. The interval between two consecutive exposures to the same odorant was 60 seconds. Note that three blocks of twenty-five trials are shown. Between each block, a 15-minute no-odor reset window was included. Hexanol was repeatedly presented during the first and third block of trials. 2-octanol was repeatedly presented during the second block of trials. A blue or a red color shade is used to differentiate the pre-stimulus period preceding the repeated hexanol or 2-octanol exposures, respectively. Gray box indicates the 4 s when the odor stimulus was presented. **(b)** The pre-stimulus spiking activities (first 15 s of each trial) of four representative projection neurons are shown (25 trials in each condition). Note that neurons have distinct pre-stimulus activity levels when a different stimulus (2-octanol) is repeatedly encountered. Further, the pre-stimulus activity levels are recovered when the same stimulus (hexanol) is repeated during different epochs. Each block is separated by a 15-minute no-odor reset window. **(c)** Comparison of total spike counts over the entire pre-stimulus window preceding repeated presentation of two different odorants (hex and 2oct). The X-axis corresponds to spike counts during the pre-stimulus period when hex was repeatedly presented. The Y-axis corresponds to spike counts during the pre-stimulus period when 2oct was repeatedly presented. Mean ± s.e.m. over 25 trials is shown for all PNs. Cells in red indicate a significant increase in spiking activity preceding 2oct exposures compared to similar epochs preceding hex exposures (P < 0.05, d.f. = 1, 48, one-way ANOVA; n = 80 PNs). Similarly, cells in blue indicate a significant decrease in spike counts for epochs preceding 2oct exposures relative to epochs preceding hex exposures, and cells in gray indicate no significant change in their baseline firing rates across the two conditions. The inset summarizes the fraction of PNs with a significant increase (red) or decrease (blue) in spike counts. **(d)** Pre-stimulus ensemble projection neuron activities were visualized using principal component analysis. The average spiking responses over the 15 s pre-stimulus window across all PNs in a particular trial (i.e. one high dimensional vector per trial) is shown as a color-coded sphere in the plot.

How do these observed changes in spontaneous activities relate to the responses evoked by the repeating stimuli? To understand this, we computed correlations between the ensemble neural activities in different time bins recorded during a single trial (baseline and odor presentation window included; **Fig 6a, b**). As can be expected, we found that the ensemble responses during the odor presentation window were highly correlated amongst themselves. No noticeable correlation between the pre-stimulus activities and the odor evoked responses were observed in the first trial. However, the ensemble activities in the pre-stimulus time window gained a negative correlation during later trials of the same odorant (the prominent ‘†’ pattern observable in the later trials). This systematic reduction in correlation between the pre-stimulus and stimulus-evoked ensemble responses was observed for all four odorants used in this study (**Fig. 6c, d**; left-most panels). Also note that the PN response immediately after the termination of the stimulus was the least correlated with the odor-evoked neural activities (**Fig. 6c, d**; middle panels), whereas the PN spiking responses during pre-stimulus and post-stimulus epochs in a given trial remained highly correlated with one another across trials (**Fig. 6c, d**; right most panels). In sum our results reveal that sensory memory about a repetitively encountered stimulus persists in the antennal lobe and generates a persistent negatively scorrelated spontaneous response, i.e. a lingering ‘negative image’ of the stimulus.

**Fig. 6:**
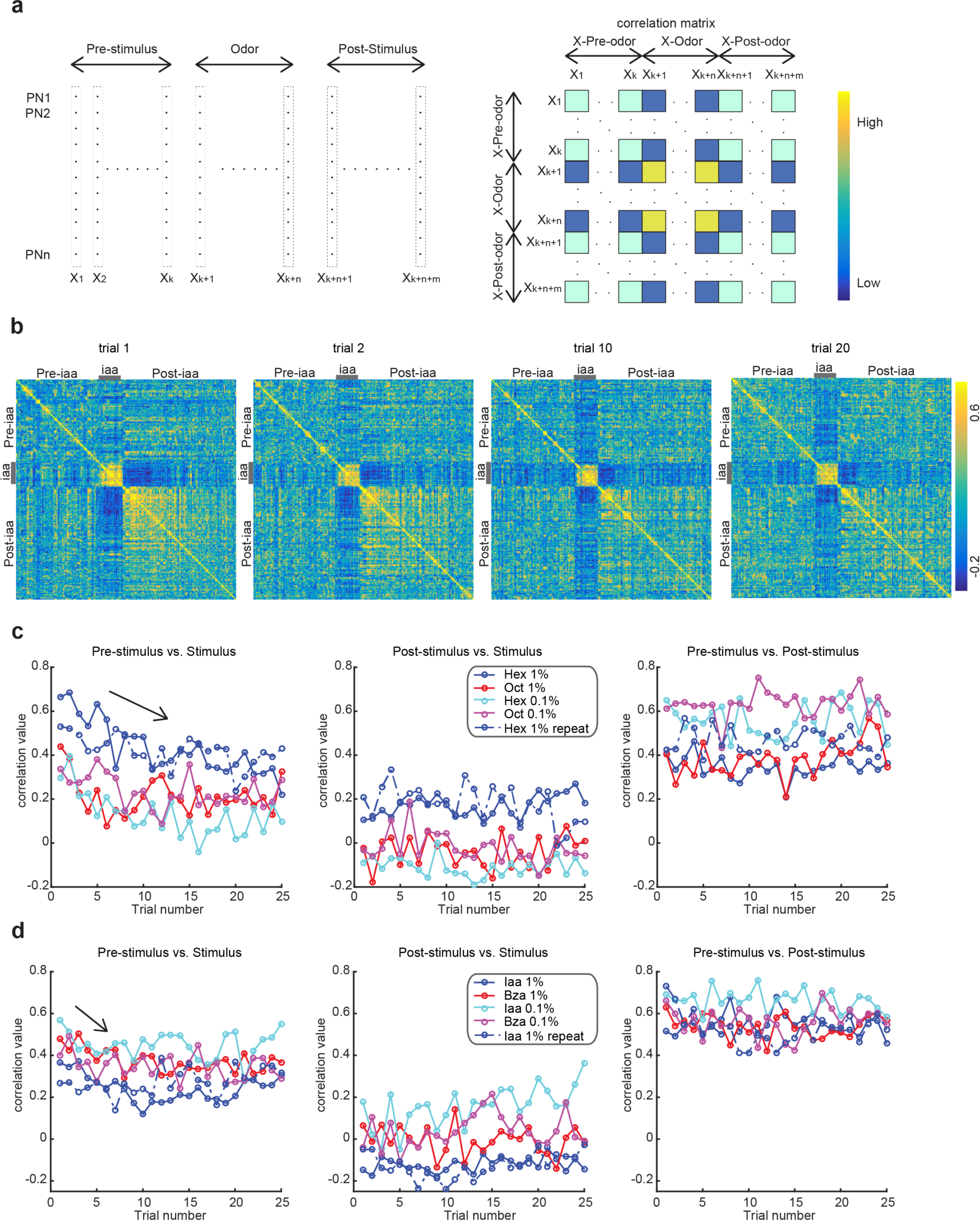
Pre-stimulus activities are negatively correlated with stimulus-evoked responses. **(a)** Schematic overview of the correlation analysis. Left panel, each rectangular column indicates a vector of ensemble projection neuron spike counts in a 50 ms time bin in one specific trial. Ensemble PN vectors during pre-stimulus, stimulus, and post-stimulus presentation periods were included in this analysis. Right panel, correlations between PN ensemble activity in a given time bin with all other epochs were computed and shown as a row in the correlation image. Diagonal pixels correspond to self-correlations. Note that the image is symmetric. **(b)** Correlation maps for different trials are shown. The 4 s stimulus presentation period is identified using a gray bar. Note that each non-diagonal pixel represents similarity between ensemble PN spike counts in one time bin versus those in another time bin. Similarly, one row or column represents the correlation between one ensemble PN activity vector with all other ensemble PN vectors. Cooler colors indicate lower correlation and hotter colors indicate higher similarity/correlations. Four representative trials are shown. Note that the correlation values between pre-stimulus PN spiking activities and odor-evoked spiking responses decrease across trials. **(c)** Left, correlations between mean pre-stimulus ensemble PN responses and stimulus-evoked population PN responses are shown when the same stimulus repeated 25 times with a 60 s inter-stimulus interval. Different colors point to different odors and concentrations; dotted line indicates results for hexanol repeat block. Middle, similar correlation plots but now showing the comparison between stimulus-evoked responses and post-stimulus activities. Right, similar plots showing correlation changes over trials between pre-stimulus and post-stimulus. **(d)** Similar plots as in **panel c** for a different set of odorant stimuli.

### Contrast-enhanced response to an unexpected stimulus

We next wondered whether the network-level adaptation allowed differential processing of repetitive vs deviant stimuli. To understand this, we examined the individual and ensemble PN responses to the deviant stimulus (**Fig. 7a**). As indicated in **Fig. 1**, the response to the repetitive stimulus diminished over trials. We observed four main PN response motifs during the catch trial (**Fig. 7b**). Responses for a subset of PNs (**cluster 1**) to the deviant stimulus was stronger during the catch trial compared to the unadapted response of the same stimulus (**Fig. 7b**; catch trial 26 in block 1 vs. trial 1 in block 2). A second cluster consisted of PNs that adapted their responses to the repetitive stimuli but recovered their responses during deviant stimulus exposure in the catch trial (**Fig. 7b – cluster 2**). A few neurons that were inhibited by target odorants maintained response suppression during the catch trial (**Fig. 7b – cluster 3**). Finally, the fourth cluster included mostly PNs that were non-responsive to both odorants (**15.52%**).

**Fig. 7:**
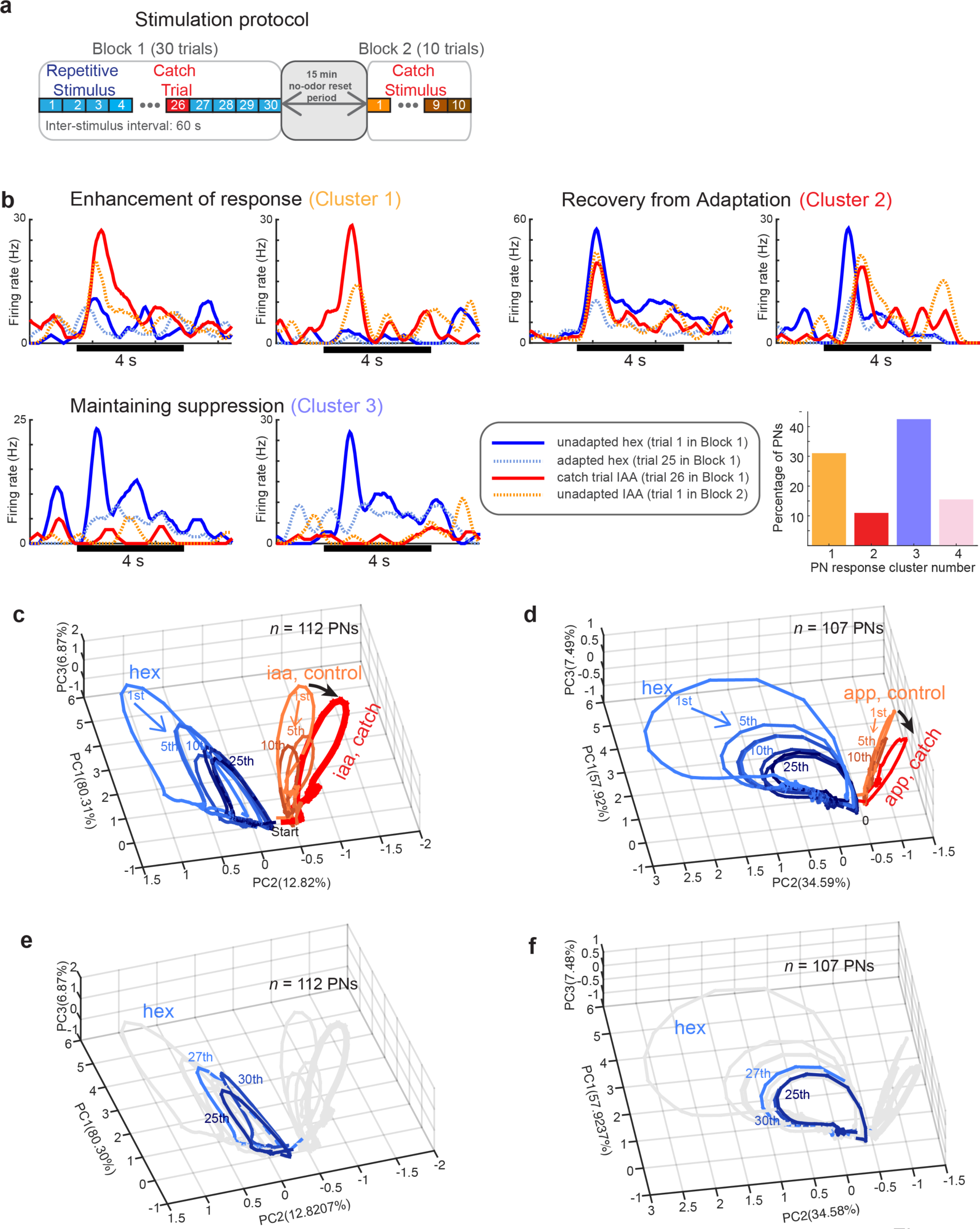
Contrast enhancement of ensemble responses to the deviant stimulus. **(a)** Two blocks of trials were used. First, a block of 30 trials where one odorant (*hex*) was presented in all trials except the 26^th^ trial (the catch trial). During the catch trial, a deviant stimulus (*iaa* or *app*) was presented. After a 15-minute no-odor reset period, a second block of ten trials of deviant stimulus was presented. This was done to determine the unadapted (first trial) and adapted (later trials) responses of the same set of PNs to the stimulus used in the catch trial. **(b)** Responses of the six representative PNs are shown as PSTHs. Four trials are compared: PN response during the first trial of the repetitive stimulus (solid blue traces), PN responses during the twenty-fifth trial of the repetitive stimulus (dashed cyan traces), PN responses during the’catch trial’ (solid red traces), and PN response to the first exposure of the deviant stimulus following the 15-minute no-odor reset period (dashed orange traces; 1^st^ trial of block 2). The PN responses types shown formed three of the main response motifs. The last category included PNs with weak or no significant response to both the repetitive and deviant stimuli. The fraction of PNs belonging to each cluster is summarized as a bar plot. **(c, d)** Similar trial-by-trial trajectory plots as in **Fig. 3**. PN ensemble response during each trial is shown as a closed loop response trajectory. Ensemble PN response to the repeat stimulus is shown in blue (hex; block 1 – trials 1 - 25). Orange response trajectories are those elicited by *iaa* or *app* during block 2 trials. The numbers next to the trajectories indicate trial number and a color gradient from light to dark is used to indicate early and late trials, respectively. The PN response trajectory elicited by *iaa* or *app* during the catch trial (block 1 – 26^th^ trial) is shown in red. The black arrows indicate the shift in the direction of PN response trajectories during the catch trial. **(e, f)** Trial-by-trial trajectory showing a modest increase in the trajectory length observed during the trial immediately following the catch trial (27^th^ trial) when the repetitive stimulus was presented again.

To understand whether these response changes were random or facilitated certain computations, we again visualized the ensemble activities elicited during each trial (**Fig. 7c, d**). Notably, while the response to the repetitive stimuli systematically diminished across trials (i.e. trajectory length shortened), the response to the deviant stimuli during the catch trial were stronger than the adapted responses elicited by the same stimuli (i.e. after the first few trials of the same stimuli immediately following no-odor reset period). Furthermore, consistent with single neuron responses (**Fig. 7b**), the neural response trajectory during the catch trial moved further away from the repetitive stimuli indicating a population-level contrast enhancement of neural representation (**Fig. 7c, d**). Responses to the repeating stimulus, post catch trial, increased modestly in response magnitude but still pattern matched well the ensemble activity evoked by that stimulus **Fig. 7e, f**).

Taken together, these results indicate that repetitive and deviant stimuli are differentially processed in this neural network. While the response to the repetitive stimuli is selectively suppressed, the ensemble neural activities are altered to emphasize the unique features of the deviant stimuli.

### Two distinct forms of short-term memory

Finally, we sought to determine what mechanisms may underlie the short-term adaptation observed. Consistent with prior findings^29,30^, we found that repeated exposure of a stimulus builds up field potential oscillations in this circuit (**Fig. 8a-c**). Power in the faster oscillation band (~20 Hz) increased rapidly over trials before converging onto a stable state. On the other hand, power in the slower oscillation band (~10 Hz) had the opposing trend and decreased over trials. Given the importance of recurrent inhibition onto projection neurons for entraining oscillations^31,32^, the strength of this negative feedback could provide a potential mechanism for olfactory habituation^33,34^. We tested this hypothesis by pharmacologically blocking fast GABAergic inhibition from local neurons onto projection neurons with picrotoxin. Bath application of picrotoxin abolished field potential oscillations (**Fig.8d, e**), but the decrease in firing rates persisted and even became exaggerated after the manipulation (**Fig. 8f-h**). These results indicate that multiple forms of short-term memory may persist in this neural network and that facilitation of inhibition does not mediate the reduction of spiking response to a repetitive stimulus.

**Fig. 8:**
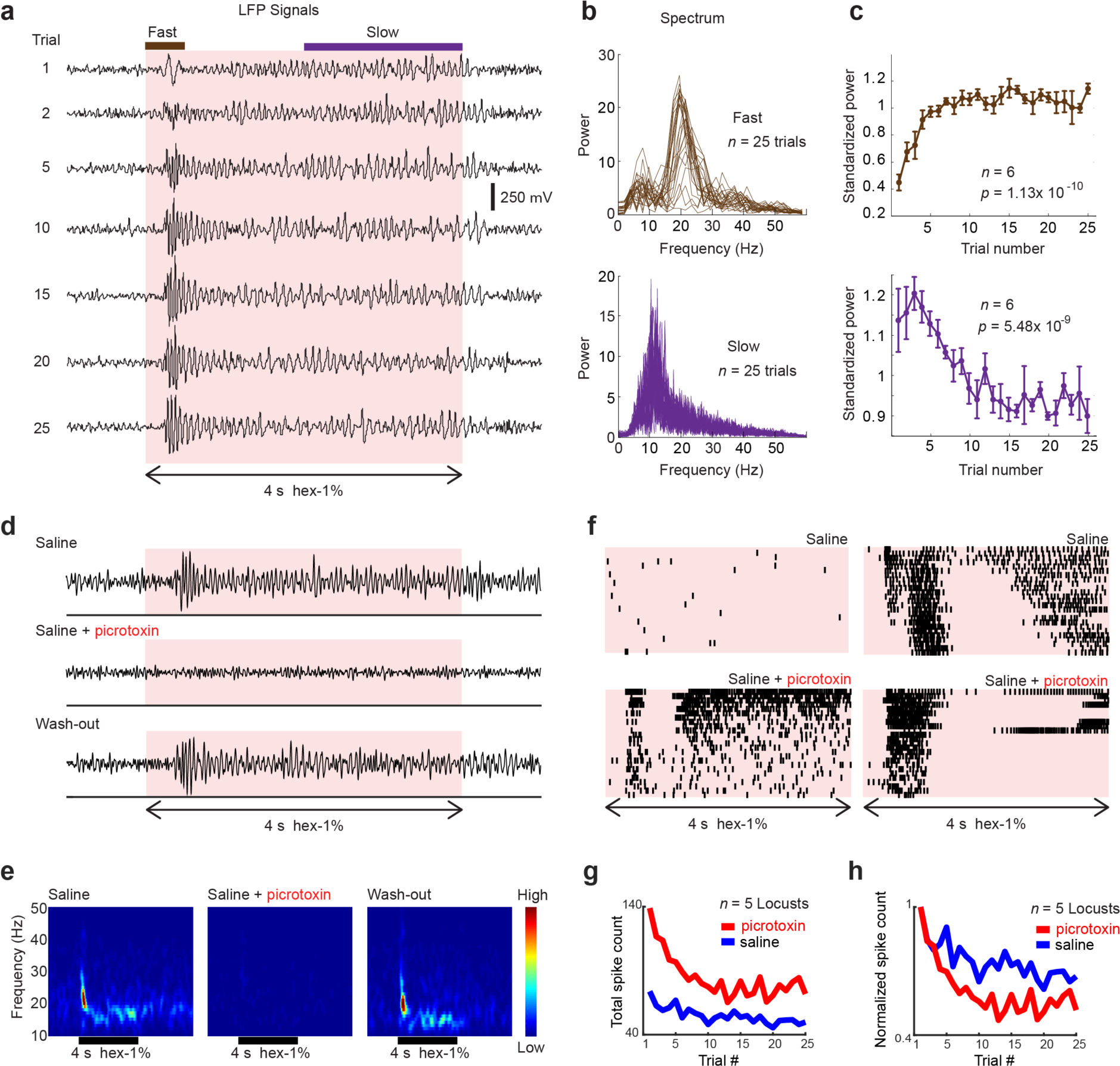
Multiple forms of short-term memory in the antennal lobe. **(a)** Local field potential responses evoked by hexanol-1% (filtered between 5 – 55 Hz) are shown for a representative subset of twenty-five consecutive trials. Two distinct analysis windows (shown on the top of the traces) are identified. Fast oscillation epoch is defined as the time segment during the first 500 ms following odor onset, while slow oscillations period refers to the duration after 2s of odor onset. The color box indicates the stimulus presentation window. **(b)** For the two time-windows identified in **panel a**, the total power in different frequencies were computed and plotted as a function of trial number. The dotted lines indicate a 20-Hz frequency band (10-30 Hz for fast oscillations; 5-25 Hz for slow oscillations) that was used for further analysis in **panel c**. **(c)** Standardized power for each trial was calculated by integrating total power over the identified frequency band and plotted. To allow comparison, the total power in the frequency band was divided by the mean response across 25 trials. *n* indicates the number of experiments. A repeated measure one-way ANOVA was used for testing the significance of the reported results. **(d)** Top, a representative LFP recording is shown. The color bar identifies a 4 s window when hexanol was presented to the locust antenna. Middle, field potential trace from the same antennal lobe after 100 µM picrotoxin bath application is shown. Bottom, LFP traces from the same location is shown after picrotoxin wash-out. **(e)** Trial averaged frequency spectrogram is shown for the 4 s hexanol puff before picrotoxin bath application (left), during picrotoxin bath application (middle), and after picrotoxin was washed out (right). **(f)** Responses of two representative PNs to 4 s hexanol puff before and during picrotoxin bath application are shown. **(g)** Summed, unsorted spike counts and normalized summed spike counts **(h)** during four second stimulus exposure period are plotted as a function of trial number. Blue and red traces show the summed spike counts before and after picrotoxin bath application, respectively.

### Discussion

The response elicited by a sensory stimulus often reduces when the same stimuli is repeatedly encountered. This form of adaptation is found in most sensory systems^26,35-38^, and is thought to allow humans and other animals to attend to other more salient or novel stimuli in their environment^1,39-41^. However, could diminishing the neural response to the recurring stimulus potentially confound other pertinent information about the same adapting stimulus such as its intensity or alter how information regarding other stimuli are transmitted? Our results indicate that adaptation does not lead to a loss of stimulus-specific information. Although neural response strength reduced with repetition, information about odorant identity and intensity were more efficiently encoded (i.e. with fewer spikes). Further, adapting to one stimulus altered how other stimuli were processed by the neural circuit. The response features that were unique to the deviant stimulus were enhanced.

Where does the memory of recently encountered stimulus reside? While olfactory sensory neuron responses have been shown to diminish upon prolonged exposures, they recover when the stimulus exposure is intermittent with large temporal gaps between consecutive exposures (**Fig. 1a**). However, the responses in the second relay center, the antennal lobe, continue to diminish even for temporally discontinuous encounters of the recurring stimulus. One reasonable explanation of these results is that the synapses between the olfactory sensory neurons and the antennal lobe PNs could depress to mediate adaptation. Our results reveal that although PN responses to the repeating odorant reduce, the spiking activity recovers to higher response levels when a deviant stimulus is presented, (**Fig. 7**, cluster 2 PNs). Therefore, it is possible that sensory input for different odorants are transmitted onto PNs through different ORN-PN synapses (i.e. multiglomerular PNs). In addition to recovery from inhibition, our results also indicate that some PNs responses to the deviant/unexpected stimulus during the catch trial was stronger than the unadapted responses evoked by the same stimulus immediately after a 15 min no-odor reset window. These latter results cannot be explained by depression of ORN-PN synapses and indicate that some aspects of this short-term memory reside within the antennal lobe neural network.

Consistent with previous findings^26^, our data also indicate that PN response changes were largest in the first few trials. However, we found that most PN spiking responses continued to change even after twenty repetitions of the same stimulus (**Fig. 2a**). While the odor-evoked PN spike trains were more consistent or stable in the later trials for a subset of PNs (**Fig. 2c**, cluster 5), the opposite was true for many others (**Fig. 2c**, cluster 1). As can be expected, compared to the individual PN responses, the ensemble response profiles remained relatively consistent across trials (**Fig. 4d, e**). As a result, although the overall response strength diminished over trials, the combination of PNs activated robustly encoded both the stimulus identity and intensity (**Fig. 3**). Intuitively, the direction of the PN ensemble response vectors changed with and robustly encoded stimulus identity and intensity. However, the length of these vectors diminished over trials (as visualized by the length of the trajectories shown in **Fig. 3b, c;** also refer **Supplementary Fig. 5a**), but did not confound information about stimulus intensity. This encoding scheme allowed adaptation-invariant representation of stimulus intensity. These results are consistent with earlier findings^20^, and extend them by including both unadapted and adapted responses (i.e. all trials) for analyses.

In addition to changes in stimulus-evoked activity, we also found that the spontaneous activity in individual PNs changed depending on the identity of the odorant that was repeatedly presented (**Fig. 5b**). The ensemble-level baseline PN responses during two non-overlapping epochs when two different odorants were presented resulted in two distinct spontaneous activity clusters (**Fig. 5d**). A prior study had reported the passive exposure to an odorant could alter the spontaneous activities such that they become more correlated with the stimulus-evoked responses. While our results also reveal that the changes in spontaneous PN spiking responses were non-random, we found that they instead became negatively correlated with the stimulus-evoked responses during that epoch (**Fig. 6**).

Notably, neural responses that are negatively correlated to a sensory stimulus (referred to as ‘negative images’) have been reported in the electrosensory lobe of electric fish^42-45^ and, more recently, in the mice auditory cortex^46-48^. In these systems, the anti-correlated responses allow cancelling sensory signals that arise from the movements generated by the animal, while allowing sensitivity to other environmental stimuli. Though functionally different, the analogy between these results with the findings of our study are hard to miss. In the antennal lobe, short-term memory results in the spontaneous activity becoming negatively correlated with the repeatedly exposed stimulus. Though the odor-evoked responses were not eliminated, they progressively became less intense, thereby allowing adaptive deemphasizing of sensory input generated by a less important stimulus.

We found that both stimulus-evoked neural activities and spontaneous activities changed in consistent manner when the same odorant was repetitively presented in different epochs (**Supplementary Fig. 2, 4**). The spontaneous ensemble PN activities in the very first trial were random. However, subsequent evolution after the first encounter with a repetitive stimulus led to reduction in correlation between neural activities observed in these two epochs (**Fig. 6**; **Supplementary Fig. 4, 5**).

Although the effect of the sensory memory was greater for the repetitive stimulus, it also altered the neural response to a deviant/unexpected stimulus (**Fig. 7**). We observed PNs whose response became less intense to the recurring odorant recovered from adaptation. More importantly, responses of PNs activated by the deviant/unexpected odor alone became stronger. Because of the latter, at a population level, the ensemble neural responses became more distinct from that of the repetitive stimulus. Note that this contrast enhancement occurs between stimuli and may potentially serve to emphasize the contrast between the deviant/unexpected stimulus with respect to the expected one.

Finally, we investigated the neural basis for this short-term memory. Prior work had shown that facilitation of recurrent inhibition onto PNs can explain behavioral habituation in fruit flies^33,34^. Further, enhancement of odor-specific inhibition was hypothesized to create a sensory memory that is a ‘negative image’ of the stimulus. This short-term memory can selectively impede transmission of a familiar stimulus^1^. Indeed, our result is largely consistent with this idea. Sensory memory of a repetitive stimulus caused the spontaneous activity in the network to progressively become negatively correlated with the odor-evoked response (**Supplementary Fig. 5b**). Further, as shown in prior studies, facilitation of inhibition was necessary and sufficient to entrain oscillatory synchronization of neural activity in the antennal lobe. However, reduction in total spiking activity persisted even when fast recurrent inhibition was pharmacologically abolished. Therefore, our results indicate multiple forms of short-term memory may co-exist in the same sensory neural network to facilitate different computations and behavioral outcomes.

## ACKNOWLEDGMENTS

We thank members of the Raman Lab (Washington University in St. Louis) for feedback on the manuscript. This research was supported by an Office of Naval Research grant (N00014-16-1-2426) and an NSF CAREER grant (#1453022) to B.R.

## AUTHOR CONTRIBUTIONS

BR conceived the study and designed the experiments/analyses. LZ and CL did all the analyses. LZ generated all the figures. DS and AC performed the electrophysiological recordings. BR wrote the paper taking inputs from all the authors and supervised all aspects of the work. LZ, AC, and DS are equally contributing first authors.

## Supplementary Methods

### Odor stimulation

For delivering odorants, we followed a protocol described in our earlier work^49^. Briefly, odorants were diluted in mineral oil to either 1% or 0.1% concentration (v/v) and sealed in glass bottles (60 ml) with an air inlet and outlet. A pneumatic picopump (WPI Inc., PV-820) was used to displace a constant volume (0.1 L/min) of the static headspace above the diluted odor-mineral oil mixture into a desiccated carrier air stream (0.75 L/min) directed toward one of the locust’s antennae. A vacuum funnel placed behind the locust preparation continuously removed the delivered odors.

The first set of experiments included multiple blocks, with twenty-five trials each, when one odorant at one intensity was repeatedly presented. Each trial in the block included a four second stimulus presentation window. The inter-stimulus interval (between trials) was sixty seconds. Different odorants at different intensities were repeatedly presented in different blocks. A 15-minute window, when no stimulus was presented, separated two consecutive blocks of trials. This window was included to reset any short-term memory that may have formed due to repeated presentation of the same stimulus^26^.

The second set of experiments involved two blocks of trials. The first block of trials included 30 trials and the second block consisted of ten trials. In the first 25 trials of block 1, hexanol at 1% v/v was repeatedly presented. In the 26^th^ trial, a puff of either isoamyl acetate 1% v/v or apple 1% v/v (a’deviant’ odorant) was presented. In trials 27-30, hexanol at 1% v/v was again presented. This was followed by fifteen-minute reset window when no stimulus was presented. In the second block of ten trials (trials 31-40), the deviant stimulus was repeatedly presented. All inter-stimulus intervals (between trials) were sixty seconds and all odor presentations were four seconds long.

### Electrophysiology

Young-adult (post-fifth instar) locusts (*Schisocerca americana*, both male and female) were used for electrophysiological experiments. First, the locusts were immobilized with both antennae intact. Then the olfactory regions of their brain were exposed, desheathed, and superfused with locust saline at room temperature. Extracellular, multiunit recordings of projection neurons (PN) were performed with a 16-channel, 4×4 silicon probe (NeuroNexus) superficially placed on the antennal lobe. Mushroom body local field potential recordings were performed using custom-made twisted-wire tetrodes (nickel-chromium wire, RO-800, Kanthal Precision Technology) placed into the mushroom body calyx. Prior to each experiment, all probes and tetrodes used were electroplated with gold to obtain impedances between 200 and 300 kΩ. Multiunit PN activity and mushroom body local field potential were recorded using a custom made 16-channel amplifier (Biology Electronics Shop; Caltech, Pasadena, CA). Signals were amplified with a 10k gain, filtered using a bandpass filter (0.3 to 6 kHz), and sampled at 15 kHz using a LabView data acquisition system. A visual demonstration of the extracellular, multiunit PN recording techniques is available online^50^.

### Electroantennogram

Electroantennogram recordings were made using intact locust antenna. Three to four distal antennal segments were cut, and the cut antennae was inserted into a micropipette filled with locust saline. An Ag/AgCl wire was inserted at the other end of the micropipette to make an electrical contact with the cut antennae. A ground electrode was inserted into the contralateral eye. The signals were acquired using a DC amplifier (Brownlee Precision). Data were collected at 15 kHz with a custom-made LabVIEW data acquisition program from 5 locusts with 25 repeated trials of 1% hexanol (4s stimulus, 60 s inter-pulse interval).

### Pharmacological Manipulation

The saline bathing the brain was displaced with locust saline containing 1mM picrotoxin. Recordings were performed after a 45-minute wait period following picrotoxin addition to the bath. Because spike-sorting is not reliable after addition of picrotoxin, total spike count was determined by including all detected peaks, without clustering. For washout experiments, standard locust saline without picrotoxin was added to the bath at a rate of 200 mL/hr for 90 minutes.

### Spike sorting

To obtain single-unit PN responses, spike sorting was done offline using the best four recording channels and conservative statistical principles^51^. Spikes belonging to single projection neurons were identified as described in an earlier work^52^. Briefly, the following criteria were used to identify single units: cluster separation > 5 noise standard deviations, number of cluster spikes within 20 ms < 6.5% of total, and spike waveform variance < 6.5 noise standard deviations. In total, 283 projection neurons (from 40 locusts) were identified.

### Neighborhood correlation analyses

To quantify the systematic changes in individual PN spike trains over repeated trials, we first binned the spiking activity in 50 ms non-overlapping time bins. This resulted in an 80-dimensional vector of spike counts during the 4 s odor presentation period in each trial. The correlation between the spike train vectors in a trial with the average spike train vector of its neighboring two trials was computed as follows:

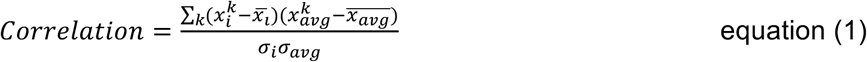

where *x*_*i*_ indicate the neural response vectors in trial i, and *x*_*avg*:_ denotes the average of two neighboring spike train vectors of trial i-1 and i+1. 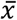 represents the mean value amongst vector components and Σ denotes the standard deviation.

For the first trial, correlation was computed with the average of the second and third trials. The last trial was correlated with the previous two trials. This metric resulted in various correlation profiles across trials for a total of 644 neuron-stimulus pairs (shown in **Fig. 2c**).

To further categorize response correlation changes across trials, we first smoothed those correlation profiles with a five-point moving average filter. Subsequently, a hierarchical clustering approach was used to identify distinct groups within similar correlation patterns across trials. A cosine similarity metric was used to calculate the pairwise distances between neurons, and a furthest neighbor clustering option was used to create the clusters shown in **Fig. 2c, d**. The elbow of a mean-squared-error of approximation was used to determine the optimal number of clusters.

### Local field potential (LFP) analysis

Raw signals were acquired at 15 kHz sampling rate. This was sub-sampled at 1 kHz and digitally filtered (5-55 Hz, 2^nd^-order Butterworth). Power spectra were obtained by fast Fourier transform of the filtered LFP, during two separate epochs (0 - 500 ms – fast oscillation window, and 2 - 4 s – slow oscillation window; all temporal window calculated from odor onset). The first-time segment revealed power spectrums centered at ~20 Hz, while the later epoch revealed power spectrums centered around 10 Hz. To allow comparison, power spectra computed for each trial were standardized by its mean response that was obtained by integrating a 20 Hz band around the peak value (Fig. 8b, between dotted lines). Repeated measures one-way ANOVA was used to determine whether there was a significant increase or decrease in power across trials.

### Tensor-based data decomposition

We first organized neural response data as a three-way array (Neuron × Time × Trials; the stimulus information was also blended into trial dimension), and then employed a direct 3-way tensor decomposition approach^53^. Here, the 3-d data cube was approximated using three loading matrices, A, B, and C with elements *a*_*if*_. (neuron dimension), *b*_*jf*_ (time dimension), and *c*_*kf*_ (trial dimension). *e*_*ijk*_ was the residual element (see the equation below). The tri-linear model was found using alternating least squares.

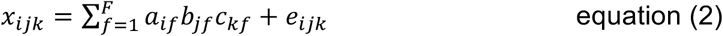

where *i, j, k* denotes the three different dimensions, and *F* indicates the total number of factors used for the analysis that was determined by the core consistency diagnostics^54^. In our case, when *F =* 3, the core consistency was above 50 %, while it dropped to below 40% when *F =* 4 (**Supplementary Fig. 1b**). Therefore, we used three factors for our data decomposition. The basis vectors *c*_*k*_ across trials in the trial dimension were shown in trial mode plots (**Fig. 3d, e**).

### Trial-to-trial odor trajectory

For this analysis, we first reconstructed the dataset by computing the outer product of the loading matrices that were obtained by the tensor decomposition. The reconstructed 3-d tensor was then unfolded into a concatenated matrix (i.e. along the trial dimension). After unfolding, the ensemble projection neuron responses were arranged as time series data of *n* dimensions (where *n* is number of neurons) and *m* steps (the number of 50 ms time bins × the number of trials). Note that only the projection neuron activities during the four second stimulus presentation window in each trial were used for this analysis. The ensemble projection neuron response vectors (in a given 50 ms time bin) were projected onto the three eigenvectors of the response covariance matrix that accounted for the most variance in the dataset, using principal component analysis. Finally, the low-dimensional points were connected in a temporal order to visualize neural response trajectories to different stimuli on a trial-to-trial basis. All trajectory plots shown in **Fig. 3** were generated after smoothing with a 3-point running average low-pass filter.

### Similarity of ensemble spiking activities and dendrogram

First, we calculated the summed spike counts during the 4 s odor presentation window for each individual projection neuron. Then the correlation similarity between two spike count profiles across projection neurons was calculated using equation 1. Similarly, in this analysis, *x*_*i*_ and *x*_*j*_ represent a *n*×1 vector (*n* = 80 for hex-2oct; *n* = 81 for iaa-bzald) for trial *i* and *j*, respectively. The dendrogram was generated by hierarchical clustering of all stimulus identities, intensities, and individual trials based on the correlation distance. The dendrogram was created in such a way that the furthest pairwise distance between any two samples assigned to an individual cluster was minimized.

### Dimensionality reduction analysis for baseline activities

For this analysis, we simply used the linear principal component analysis. We averaged the 15 s pre-stimulus spiking activities for each projection neuron in a given trial. This resulted in a n-dimensional vector for each trial (n = number of projection neurons recorded). To compare baseline projection neuron activities during different pre-stimulus epochs, we concatenated projection neuron firing count vectors during these epochs and performed a linear principal component analysis. The high-dimensional baseline PN activity vectors in each trial were projected along the leading three eigenvectors of the data covariance matrix.

### Correlation analysis between baseline and odor-evoked response

Both the pre-stimulus and stimulus-evoked activities were binned in 200 ms non-overlapping time bins for each individual trial to obtain population neuron response vectors as shown in **Fig. 6a**. Correlation value between two responses vectors then were calculated using equation 1 (here *x*_*i*_ represents population neuron response vectors in i^th^ time bin). Each pixel in correlation plot (**Fig. 6b)** indicates the correlation value between two projection neuron spike count vectors. Finally, we used correlation values between mean firing rates during different periods to obtain correlation values across trials (as shown in **Fig. 6c**). The mean pre-stimulus ensemble PN vector was obtained by taking the mean spiking activities across the entire 15 s pre-stimulus period. The mean stimulus-evoked population PN response was obtained by taking the mean of the entire 4 s stimulus-evoked responses. The mean post-stimulus activity vector was computed by averaging PN responses in a 4 s window starting after 1 s of stimulus termination.

### Clustering analysis of firing rates profile across trials (Fig. 7)

For each neuron, we calculated the mean firing rates during 4 s odor presentation for each trial and concatenated them into a 40×1 vector (30 trials from block 1 and 10 trials from block 2). This resulted in mean firing rates profiles across trials for a total of 219 neuron-odor pairs. Then, we performed a hierarchical clustering analysis using correlation distance to obtain four distinct response motifs. The correlation distance between two mean firing rates profiles was calculated as follows (here *x*_*i*_ and *x*_*j*_ represents 40×1 vector for i^th^ and j^th^ neuron-stimulus pair).

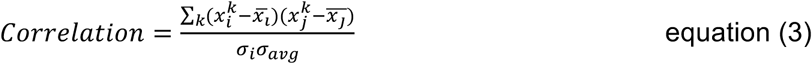

### Justification of statistical tests

All statistical significance tests done in the manuscript were two sided. No statistical methods were used to predetermine sample sizes, but our sample sizes are similar to those reported in previous publications in the field^19,24^.

Two-way ANOVA test was used to compare the means of columns and rows of the observation in a two-dimensional dataset. For **Fig. 2d**, the data in different rows represented changes in stimulus conditions, while data in different columns represented changes in cluster numbers. Similarly, in the analysis of **Fig. 3a**, the row factor represented different odor intensities, while column factor indicated different trial numbers. The observations for within and between groups were assumed to be independent.

Repeated measures one-way ANOVA was used to assess the significant changes of the mean LFP oscillatory power across repeated trials (**Fig. 8b, c**). The measurement was repeated on the same sets of the insects, and data in each group satisfied the normality (Jarque-Bera test) and equal variance assumption (Levene’s test).

---

**Supplementary Figure 1:**
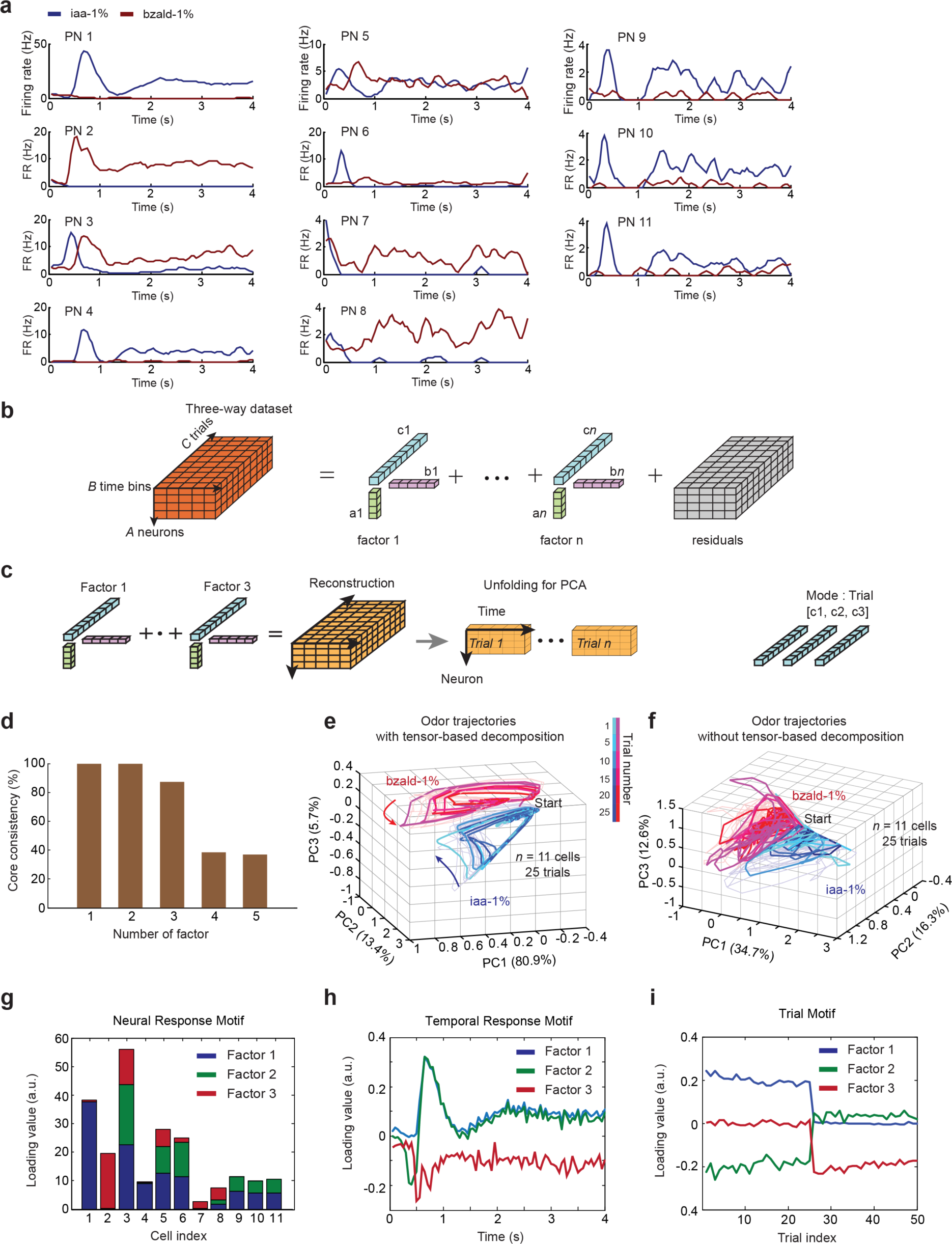
Tensor-based data decomposition for trial-by-trial analysis of ensemble neural response profiles. **(a)** Peristimulus-time histograms (PSTH) are shown for 11 simultaneously collected PNs from a single locust antennal lobe. Responses to two odorants at a single concentration are shown: iaa (1% v/v dilution) and bzald (1% v/v dilution). Only firing rate responses during the four seconds of odor pulse duration are shown for each neuron. **(b)** A three-dimension neural data array can be approximated as a sum of multiple rank-1 tensors that are obtained through outer product of three loading vectors a_f_ (neuron dimension), b_f_ (time dimension), and c_f_ (trial dimension) (subscript f ranges from 1 to the total number of factors). **(c)** The loading vectors are used to reconstruct the three-way data. This reconstructed data cube is unfolded and concatenated with respect to the trial dimension. Subsequently, a dimensionality reduction technique (e.g. PCA) is applied for odor response trajectory visualization. **(d)** A core consistency diagnostic was used to determine the number of factors to be used in the tensor-based decomposition analysis. Here, the core consistency metric was above 80% for up to three factors decompositions, but fell below 40% when the factor number was increased to four or greater. Therefore, a three-factor decomposition was used for all tensor reconstructions in this study. **(e, f)** Odor-evoked ensemble neural spiking activities from a single locust after dimensionality reduction using principle component analysis are shown (PCA). Each axis corresponds to one of the first three principal components that capture a certain amount of variance in the dataset. The odor trajectories generated by iaa and bzald are shown in blue and red, respectively for all 25 trials. For those of trial 1, 5, 10, 15, 20, and 25, neural trajectories are highlighted with color gradient that goes from light to dark. All trajectories start from a pre-stimulus baseline that is labeled ‘Start’. Arrows indicate the direction of response evolution over time. The PCA following the tensor-based decomposition method were used to generate odor trajectories in panels **e,** while a direct unfold-then-PCA approach was used to generate results in panel **f**. **(g, h, i)** Loading vectors obtained from the tensor-based decomposition corresponding to the neuron (**g**), time (**h**), and trial (**i**) dimensions are shown.

**Supplementary Figure 2:**
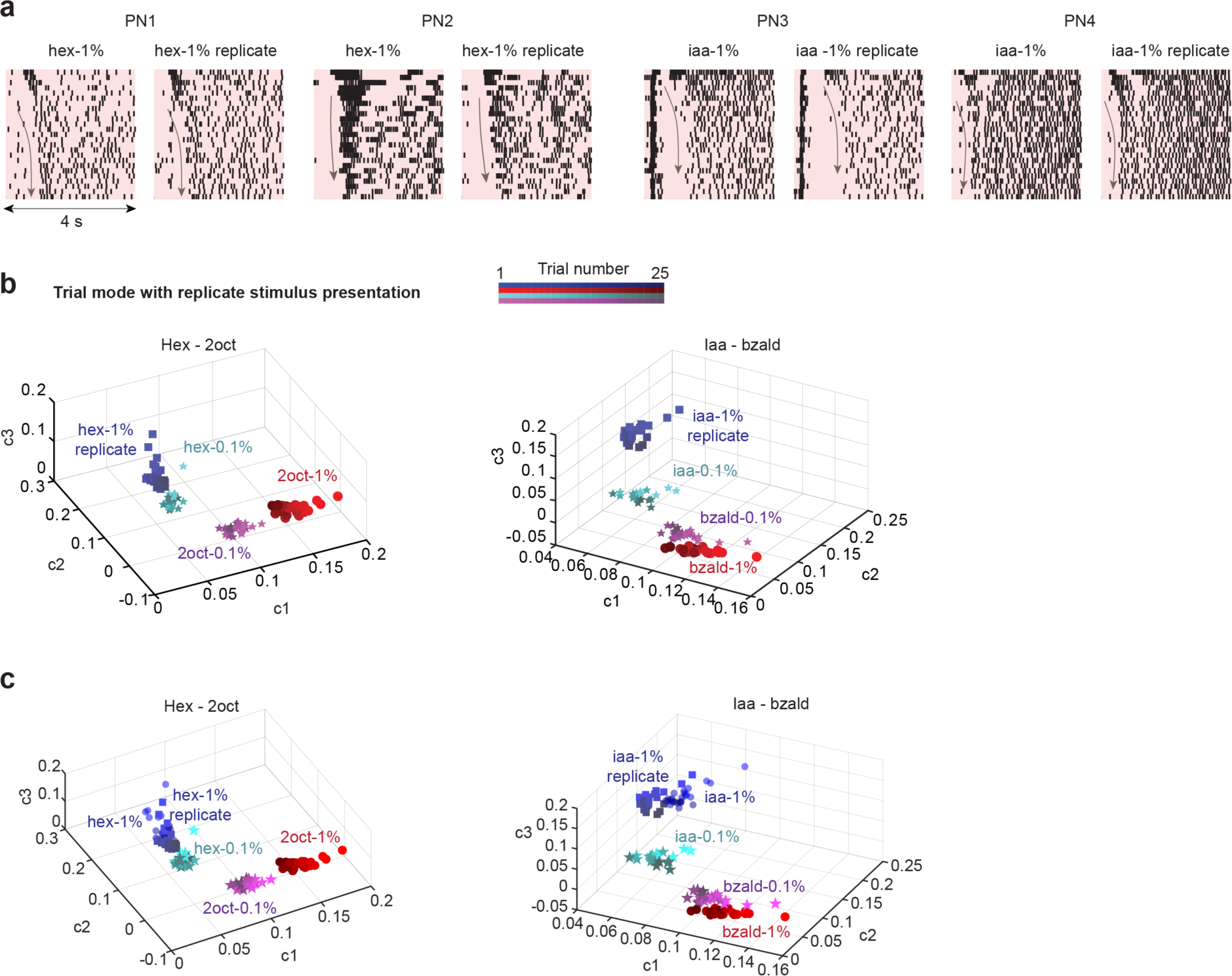
Trial-by-trial response changes are reproducible. **(a)** Raster plots of four representative PNs responses to two blocks of trials when the same stimulus was presented repeatedly in all 25 trials. A no-odor reset period of greater than 15 minutes preceded each block of trials. The color box represents 4-s stimulus duration, and the arrows in the box indicate the systematic change in individual PN responses. **(b)** Similar plots as in **Fig. 3d, e** are shown for the repeated block of twenty-five trials (hex 1% or iaa 1%) used in this study. Lower concentrations of the same stimulus are shown to allow qualitative comparisons with panels shown in **Fig. 3d, e**.

**Supplementary Figure 3:**
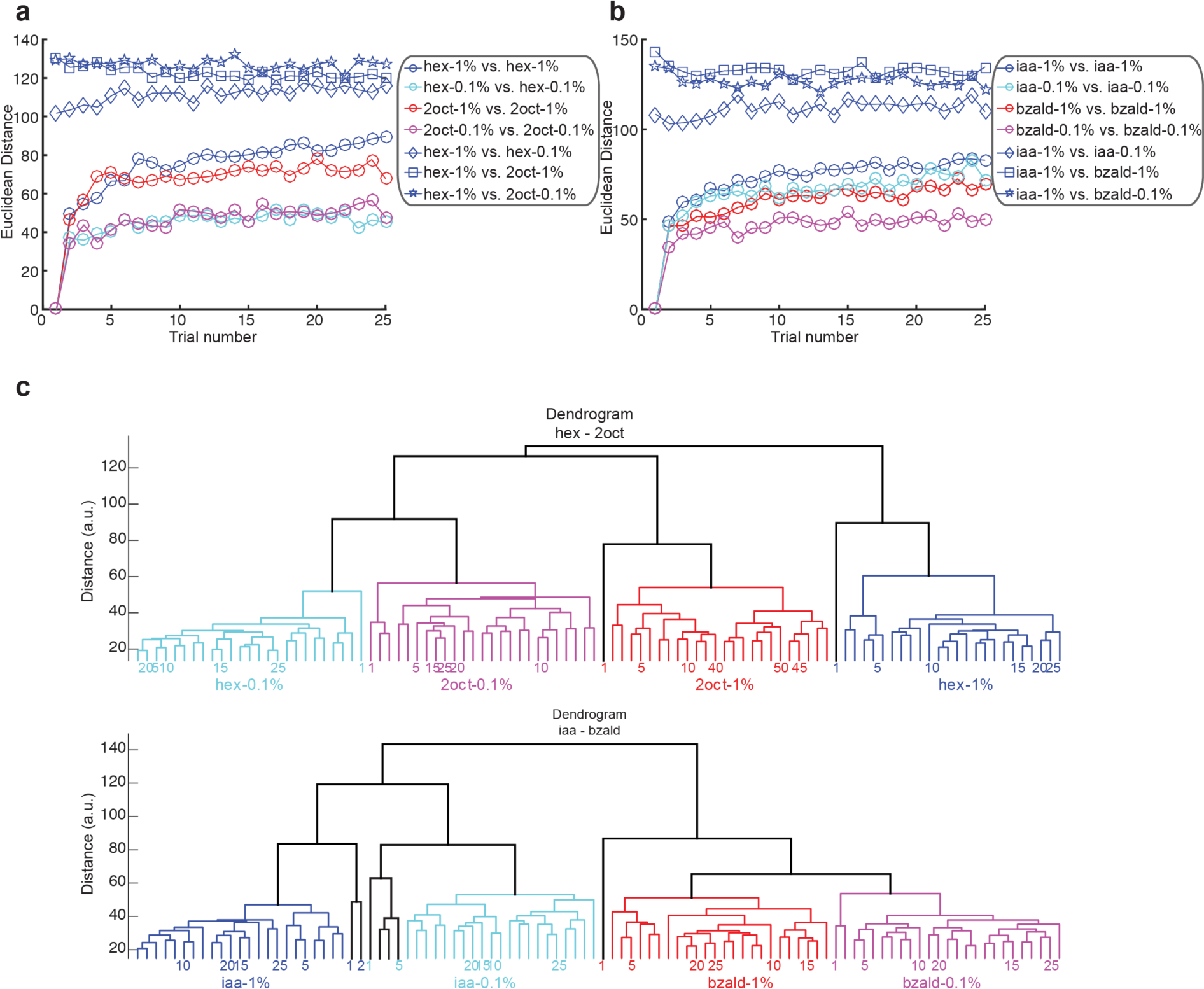
Euclidean distance based analysis of odor identity and intensity. **(a)** A comparison of combinatorial PN response profiles activated by the same odorant across trials (‘ ο’), same odorant across intensities (‘ ◊’), between the different odorants presented at same dilution levels (‘ ◻’) and between different stimuli presented at different intensity levels (‘ ★’) are shown as a function of trial number. Note that compared to results shown in **Fig. 4d, e** where a correlation distance metric was used, here we used a Euclidean distance. Comparisons are made with respect to the first trial response of that odorant. For comparisons between odorants and across intensities levels, Euclidean distance with respect to the first trial of hexanol at 1% was computed and plotted. **(b)** Similar plot as in **panel a**, but analyzing responses of a different set of PNs to a different pair of odorants (iaa and bzald). For comparisons between odorants and intensities levels, similarity with respect to the first trial of iaa at 1% was computed and plotted. **(c)** A dendrogram was generated using a Euclidean distance metric comparing trial-by-trial ensemble spiking activities evoked by two different stimuli at two different intensities. Two major response clusters that correspond to stimulus identity and intensity were identified. The number at the leaf node represents the trial number. However, note that for the hexanol and 2-octanol case, the grouping is first by intensity and then by identity. Also, the early trials of all odorants (particularly trials one and two) form a separate sub-cluster.

**Supplementary Figure 4:**
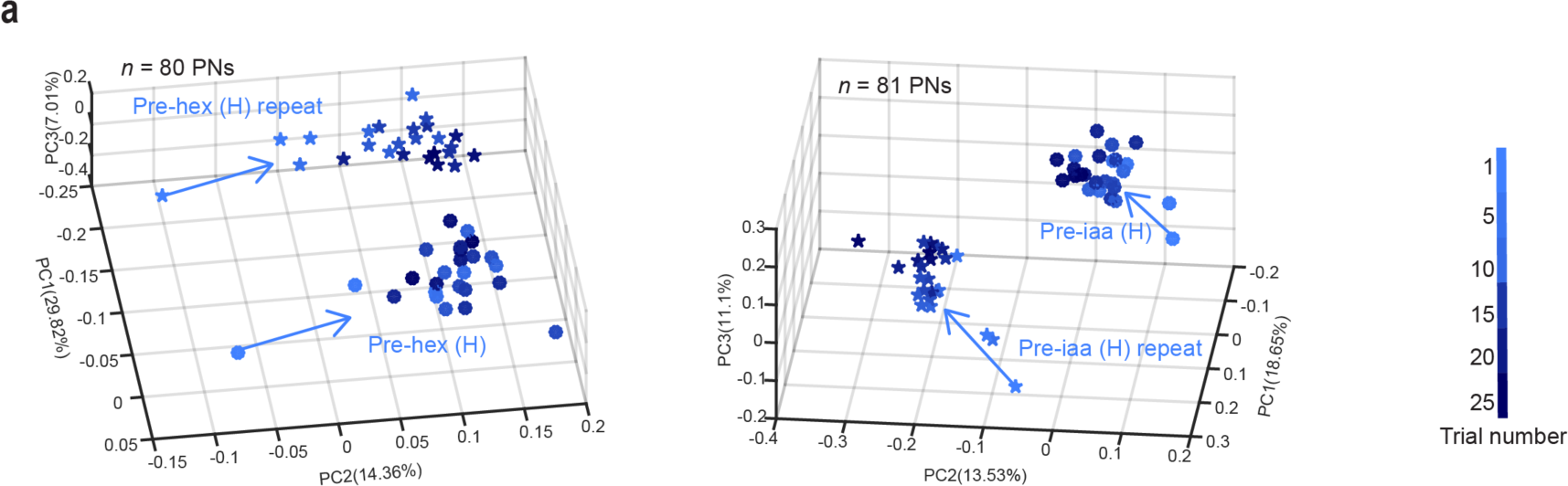
Sensory memory regarding the repeated stimulus persists in the baseline response. Similar plot as **Fig. 5d**, but pre-stimulus ensemble projection neuron activities when the same stimulus is repeated during two different epochs are visualized. The direction in which the baseline activities change in between stimulus exposures is highly reproducible when the same stimulus is repeated in different time segments.

**Supplementary Figure 5:**
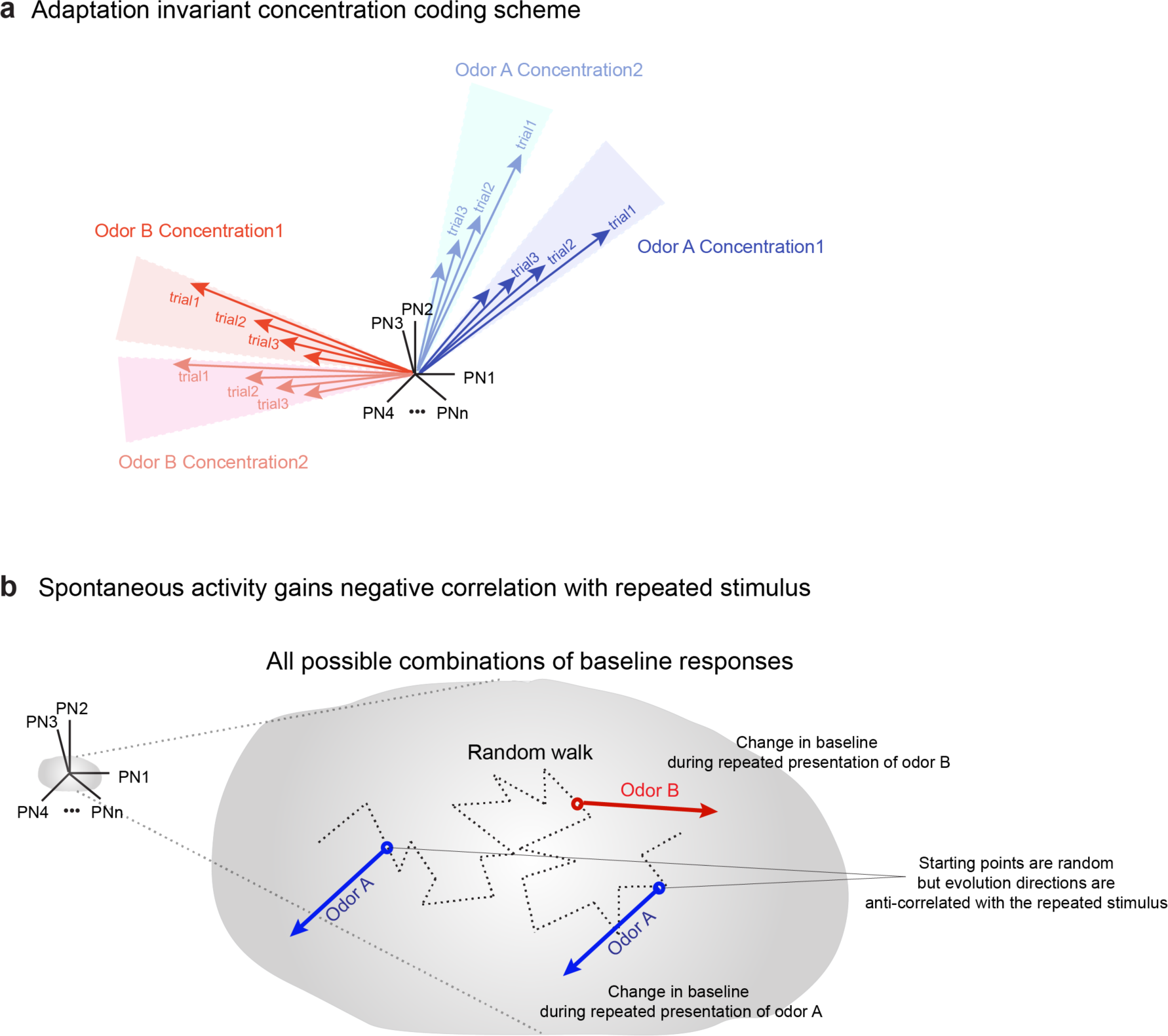
Schematic representation of the main ideas of the manuscript. **(a)** A schematic showing how odor identity, concentration can be encoded in an adaptation invariant manner. The direction of the high-dimensional vector is determined by the combination of the neurons activated and the response strength determines the length of the vector. Our results indicate that the ensemble of PNs activated changes subtly with stimulus intensity and drastically with stimulus identity. Although the response strength reduces upon repetition, the combination of PNs activated is maintained. **(b)** A schematic showing how baseline ensemble neural responses changes upon repeated stimulus exposure. Without any odor stimulus, there is some inherent variability in the ensemble PN baseline response which can be represented as a random walk in the high dimensional state space (dotted line). When an odor stimulus is encountered repeatedly, ensemble PN baseline changes in a specific direction in the state space. The observed changes in the baseline activity are stimulus specific (red and blue arrows). However, note that repeated encounters of the same stimulus results in similar shifts in baseline but the starting points in the state space are different (indicated using two blue arrows but with different starting points).

